# Fragmentation Through Polymerization (FTP): A New Method to Fragment DNA for Next-Generation Sequencing

**DOI:** 10.1101/505966

**Authors:** Konstantin B. Ignatov, Konstantin A. Blagodatskikh, Dmitry S. Shcherbo, Tatiana V. Kramarova, Yulia A. Monakhova, Vladimir M. Kramarov

## Abstract

Fragmentation of DNA is the first and very important step in preparing nucleic acids for NGS. Here we report a novel Fragmentation Through Polymerization (FTP) technique, which is simple, robust and low-cost enzymatic method of fragmentation. This method generates double-stranded DNA fragments that are suitable for the direct use in NGS library construction, and allows to eliminate the need of an additional step of reparation of DNA ends.

## Introduction

Next Generation Sequencing (NGS) has become one of the major and widely used techniques in genomic research and genetic diagnostics. Fragmentation of DNA is the first main step in preparing sequencing library for NGS. The well-known NGS technologies, like Illumina or Ion Torrent, generate a plethora of reads with lengths under 600 - 1000 bases. The quality of NGS is largely dependent on the quality of the DNA fragmentation, thus making this step utterly critical in the process of library construction.

There are three main approaches to shorten long DNA for the library preparation: physical (by using acoustic sonication or by hydrodynamic shearing), enzymatic (based on the usage of endonucleases or Transposase) and chemical shearing (utilizing hydrolyze of DNA at heating with divalent metal cations) [1, 2].

Acoustic shearing with Covaris ultrasonicators (Covaris, Woburn, MA, USA) is currently the gold standard for fragmentation at random nucleotide locations for a NGS library construction, but it requires a significant upfront capital investment and can be financially inaccessible for many laboratories [3].

Enzymatic methods versus acoustic shearing have a similar efficiency and do not need expensive equipment [2]. Commercially available Fragmentase (New England Biolabs, Ipswich MA, USA) and Nextera tagmentation (Illumina, San Diego, CA, USA) are the most popular enzymatic techniques. Nextera uses a transposase to simultaneously fragment and insert adapters onto dsDNA [4]. Fragmentase contains two enzymes: one randomly nicks dsDNA and the other cuts the strand opposite to the nicks [2].

DNA fragments obtained by physical fragmentation or by Fragmentase method require a reparation of DNA ends for the following ligation with adapters during NGS library construction [1, 2]. To reduce a reparation stage and improve a protocol of NGS library generation, we have developed a new enzymatic method for DNA fragmentation: Fragmentation Through Polymerization (FTP). Our FTP method is based on the use of two enzymes: non-specific endonuclease, which randomly nicks dsDNA (DNase I) and thermostable DNA polymerase with strong strand-displacement activity (SD DNA polymerase [5]). At the first stage of FTP DNase I introduces nicks into dsDNA, and at the second stage SD polymerase elongates 3’-ends of the nicks in a strand-displacement manner. As a result, FTP generates a number of double-stranded DNA fragments (Fig. 1) that are suitable for direct ligation with adapters (without a reparation of the ends).

**Fig 1.**
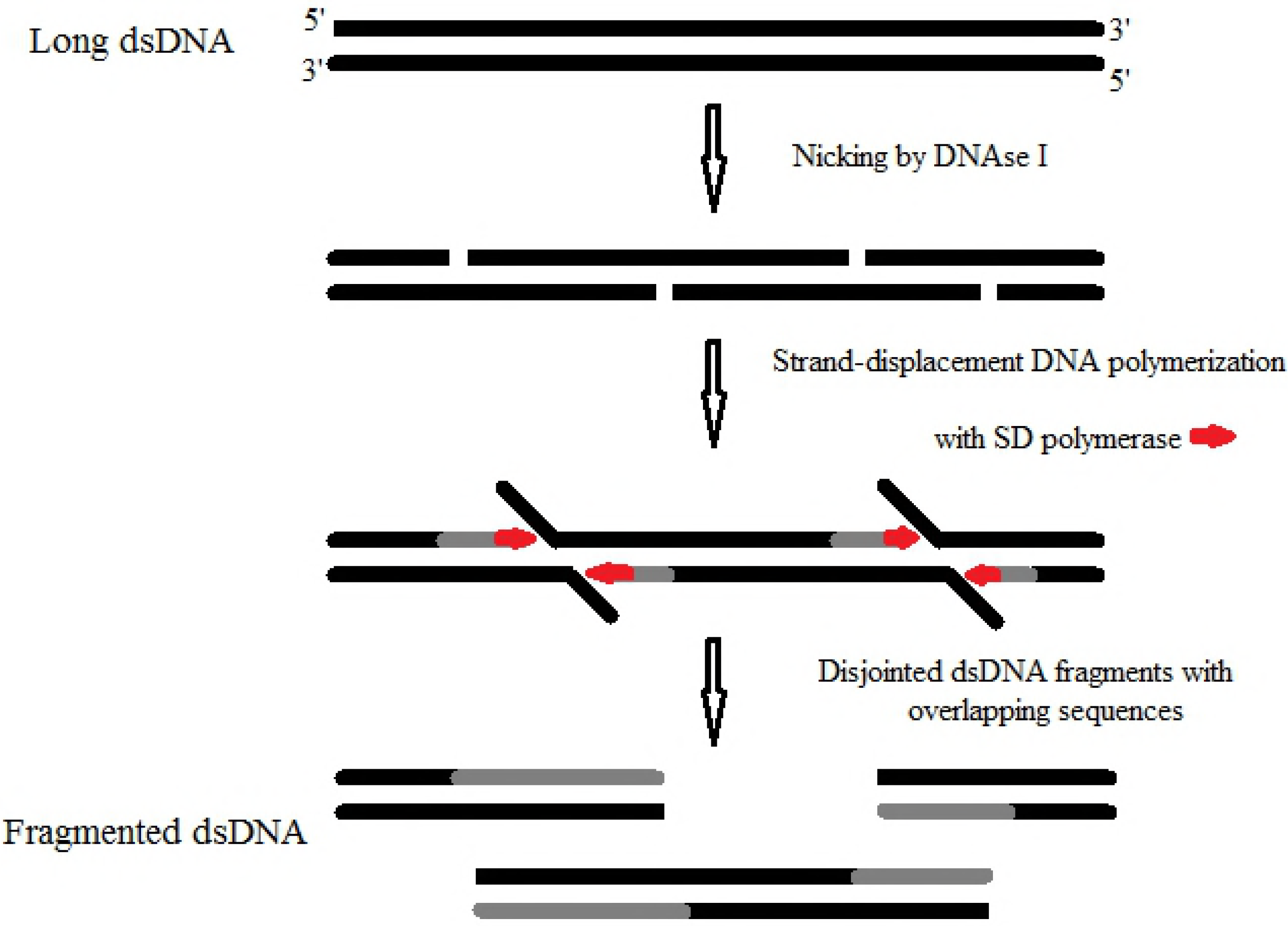
General overview of dsDNA Fragmentation Through Polymerization (FTP) method. FTP method is based on two enzymatic reactions: a DNA nicking with DNase I and strand-displacement DNA polymerization with SD DNA polymerase. As a result, a number of double-stranded DNA fragments with overlapping sequences are generated. *De novo* synthesized DNA is indicated in grey, and SD polymerase is indicated in red.

Here we describe the detailed FTP method of DNA fragmentation and compare it with the well-known and widely used Fragmentase technique (New England Biolabs). Systematic comparison of Fragmentase with other fragmentation methods has been described earlier [2].

## Materials and methods

### Enzymes and reagents

Lyophilized DNase I (deoxyribonuclease I from *Bovine pancreas*) was obtained from Sigma-Aldrich (St Louis, MO, USA) and solved in the storage buffer (50% glycerol, 100 mM NaCl, 0.2 mg/ml BSA, 1 mM EDTA, 0.2 mM DTT, 20 mM Tris-HCl, pH=8.0) up to 1 mg/ml.

SD DNA polymerase (50 U/µl) and the reaction buffer were supplied by Bioron GmbH, (Ludwigshafen, Germany). *E.coli* BL21(DE3) gDNA was supplied by Evrogen JSC (Moscow, Russia). dNTPs were obtained from Bioline Limited (London, GB).

NEBNext dsDNA Fragmentase and NEBNext Ultra II DNA Library Prep kit were supplied by New England Biolabs, Inc. (Ipswich, MA, USA).

### dsDNA Fragmentation Through Polymerization (FTP)

For fragmentation, 200 ng gDNA of *E.coli* strain BL21(DE3) was added to the reaction mixture: 1X reaction buffer for SD polymerase (Bioron GmbH), 3.5 mM MgCl_2_, dNTPs 0.25 mM (each), DNase I 1.5 ng/µl, SD DNA polymerase 1.5 U/µl. The total volume of the reaction was 25 µl. The reaction mixture was completed at 4°C (wet ice). The fragmentation of gDNA was carried out by two-step incubation: 20 minutes at 30°C and then 20 minutes at 70°C. For incubation we used thermal cycler with heated lid. The reaction was stopped by cooling down to 10°C. The mixture was diluted 1:1 with sterile water and fragmented DNA was purified with SPRI beads.

### DNA Fragmentation with NEBNext dsDNA Fragmentase

gDNA of *E.coli* strain BL21(DE3) was digested by using NEBNext dsDNA Fragmentase (New England Biolabs, Inc.), following the manufacturer’s protocol. Briefly, 200 ng of gDNA were added to the reaction mixture (total volume 25 µl):1X Fragmentase Reaction Buffer v2, 10 mM MgCl, and 1X dsDNA Fragmentase. The mixture was incubated at 37°C for 20 minutes. The digestion was stopped by adding EDTA up to 100 mM. The mixture was diluted 1:1 with sterile water and fragmented DNA was purified with SPRI beads.

### Preparation of NGS libraries

We prepared four NGS libraries from four different samples of Fragmentase-digested gDNA and four NGS libraries from four different samples of FTP-digested gDNA. NGS libraries were generated using NEBNext Ultra II DNA Library Prep kit (New England Biolabs, Inc.) according to the manufacturer’s instruction. The conventional procedure for Fragmentase digested DNA included: repair of DNA ends with “NEBNext Ultra II End Prep Enzyme Mix”, addition of adapters to the DNA fragments by “NEBNext Ultra II Ligation Master Mix” and amplification of the adaptor-ligated DNA fragments with “NEBNext Ultra II Q5 Master Mix”. Input amount of each DNA sample was 200 ng. The library indexing and amplification were performed for 5 PCR cycles as described in the kit’s manual.

NGS libraries from FTP digested gDNA were constructed by NEBNext Ultra II DNA Library Prep Kit procedure, but with the exception of DNA end reparation stage.

After the amplification stage, all libraries were quantified with Quant-iT PicoGreen dsDNA Assay Kit (Molecular Probes, Inc., Eugene, OR, USA), pooled (500 ng of each) and purified with AMPure XP beads.

### Next Generation Sequencing (NGS) and bioinformatic analysis

The pooled libraries were sequenced on the Illumina MiSeq Instrument (Illumina, California, USA) with a 300 cycles MiSeq Sequencing Kit v2, paired-end mode, resulting in 12×10^6^ reads. Each of the reads was ∼150 nt long. The FASTQ files generated on the instrument were uploaded to NCBI SRArchive under project ID: PRJNA509202.

The FASTQ files were quality controlled using FASTQC v0.11.4 (Babraham bioinformatics, Cambridge, UK). PHRED scores were calculated by FASTQC v0.11.4. Adapters were trimmed with FLEXBAR v.2.5 [6]. Filtered reads with a minimum length of 30 bp were subsequently aligned to the *E.coli* BL21(DE3) genome (NCBI Reference Sequence: NC_012971.2) using BOWTIE2 software v2.3.4 [7]. Random samples of reads were generated using Seqtk software (https://github.com/lh3/seqtk). *De novo* assembly of contigs was carried out with SPAdes tool v3.10.1 (http://cab.spbu.ru/software/spades/). Statistics were calculated using QUAST software v5 [8, 9] (http://quast.sourceforge.net/).

## Results and discussion

### Digestion of gDNA by Fragmentation Through Polymerization (FTP) method

We compared two enzymatic methods of dsDNA fragmentation for NGS library construction: digestion with Fragmentase from New England Biolabs and Fragmentation Through Polymerization (FTP). FTP method consists of two consequent enzymatic reactions: random DNA nicking and elongation in a strand-displacement manner of the 3’-ends of nicked DNA. As a result, a number of double-stranded DNA fragments with overlapping sequences are generated. The general overview of FTP method is outlined in Figure 1.

We carried out FTP in one-tube format as described in the “Materials and methods”. Mesophilic DNase I and thermophilic SD DNA polymerase were added to the reaction mixture that contained gDNA of *E. coli* strain BL21(DE3). The reaction was incubated at 30°C for 20 minutes, plus additional 20 minutes at 70°C. DNase I has the optimum performance temperature between 30°C and 40°C. During the first stage of incubation at 30°C, DNase I introduced nicks into the dsDNA. In order to optimize an average size of the obtained fragments we tested different DNase I concentrations and/or incubation times (results are not shown). During the second stage, the DNase I was heat-inactivated and the SD polymerase was activated by increasing the reaction temperature up to 70°C. The SD polymerase is the Taq DNA polymerase mutant that has a strong strand displacement activity and high thermostability (up to 93°C) with optimum of enzymatic activity at 70-75°C [5]. These properties of SD DNA polymerase, in combination with the robust polymerase activity, make it very suitable for the application in FTP technique. In summary, DNase I generated 3’-ends by nicking dsDNA at 30°C, followed by SD polymerase that used these ends for strand displacement DNA polymerization at 70°C and subsequently disjointed dsDNA fragments. As a result, the fragments with an average size about 500 bp (in a range 150 – 1500 bp) were obtained from the intact gDNA. Agarose-gel electrophoresis of gDNA fragmented by FTP is demonstrated in Figure 2. As it is seen, a cooperative work of DNase I and SD polymerase is required for the perfect DNA fragmentation (Fig. 2, lanes 4, 5).

**Fig 2.**
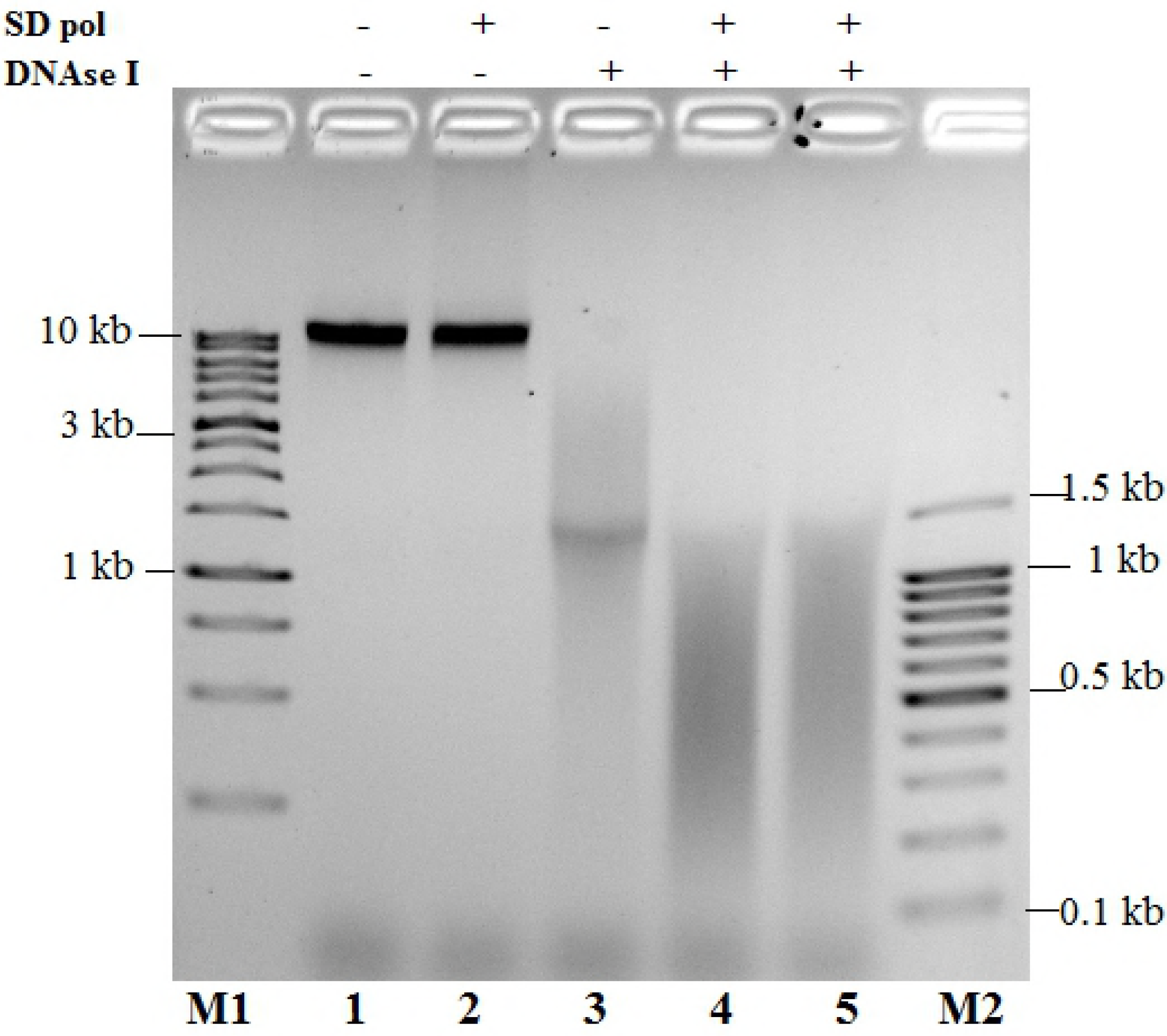
Agarose-gel electrophoresis of gDNA fragmented by FTP method. gDNA of *E.coli* BL21 was incubated as described in the “Materials and Methods”: without enzymes (lane 1), with SD polymerase (lane 2), with DNase I (lane 3), and with both DNase I and SD polymerase (lane 4, 5). **M1** – 1 kb DNA Ladder; **M2** – 100 bp DNA Ladder.

Fragmentase and other methods of fragmentation (with the exception of Illumina’s Nextera tagmentation) generate DNA fragments by introducing nicks and counter nicks in DNA strands that disassociate at 8-12 nucleotides downstream or upstream from the nick site. Thus, the generated fragments need a repair of DNA ends for the following NGS library construction [1, 2]. Unlike in other methods, in FTP the DNA fragments are separated by strand-displacement DNA polymerization and not by counter nicks (Fig. 2 demonstrates that SD polymerase is required for the fragment disassociation). As a result of FTP, double-stranded DNA fragments have ends that are suitable for direct NGS library construction and an additional step of DNA ends reparation is no longer necessary.

### NGS library constructions from Fragmentase and FTP digested gDNA

Two techniques, FTP and standard Fragmentase, were used to digest the gDNA of *E. coli* strain BL21(DE3). The fragmented DNA samples were then used for the construction of NGS libraries with NEBNext Ultra II DNA Library Prep Kit from New England Biolabs. Four libraries were prepared from the DNA samples digested with Fragmentase by the standard protocol, which included the stage of DNA end repair.

Another four libraries were prepared using the same NEBNext kit, but the DNA samples for these libraries were generated by FTP method, without the stage of DNA reparation. It is worth noting that when the DNA fragments are obtained by physical fragmentation or by Fragmentase method, the reparation of the DNA ends is required for the library construction [1, 2]. The FTP method does not require this step, therefore, the procedure of NGS library preparation is more simple.

The DNA amount in each library was quantified with Quant-iT PicoGreen dsDNA Assay Kit. All libraries generated with both Fragmentase and FTP method contained similar amounts of ds DNA, about 800 ng. This result shows that the efficiency of NGS libraries generation with FTP method is comparable to the efficiency of NGS library generation with Fragmentase technique.

### Assessment of NGS libraries generated from Fragmentase and FTP digested gDNA by Next Generation Sequencing

The obtained NGS libraries of gDNA *E.coli* BL21(DE3) were sequenced at 48× depth with an Illumina MiSeq Instrument. The raw data (about 220 Mb for each DNA sample) generated in this study have been deposited in the National Centre for Biotechnology Information (NCBI) Sequence Read Archive under BioProject accession number PRJNA509202 (https://www.ncbi.nlm.nih.gov/sra/PRJNA509202).

Different fragmentation and NGS library preparation protocols could potentially affect quality of reads. We therefore estimated the quality of reads as described in [2] for comparison of different fragmentation methods. PHRED quality scores for each base provide a sequencing error estimate and are hence a good tool to assess the quality of sequences and to compare the reliability of different sequencing runs on the same instrument [10]. We did not detect any significant differences in the quality scores obtained from the Fragmentase and FTP NGS libraries (Fig. 3).

**Fig 3.**
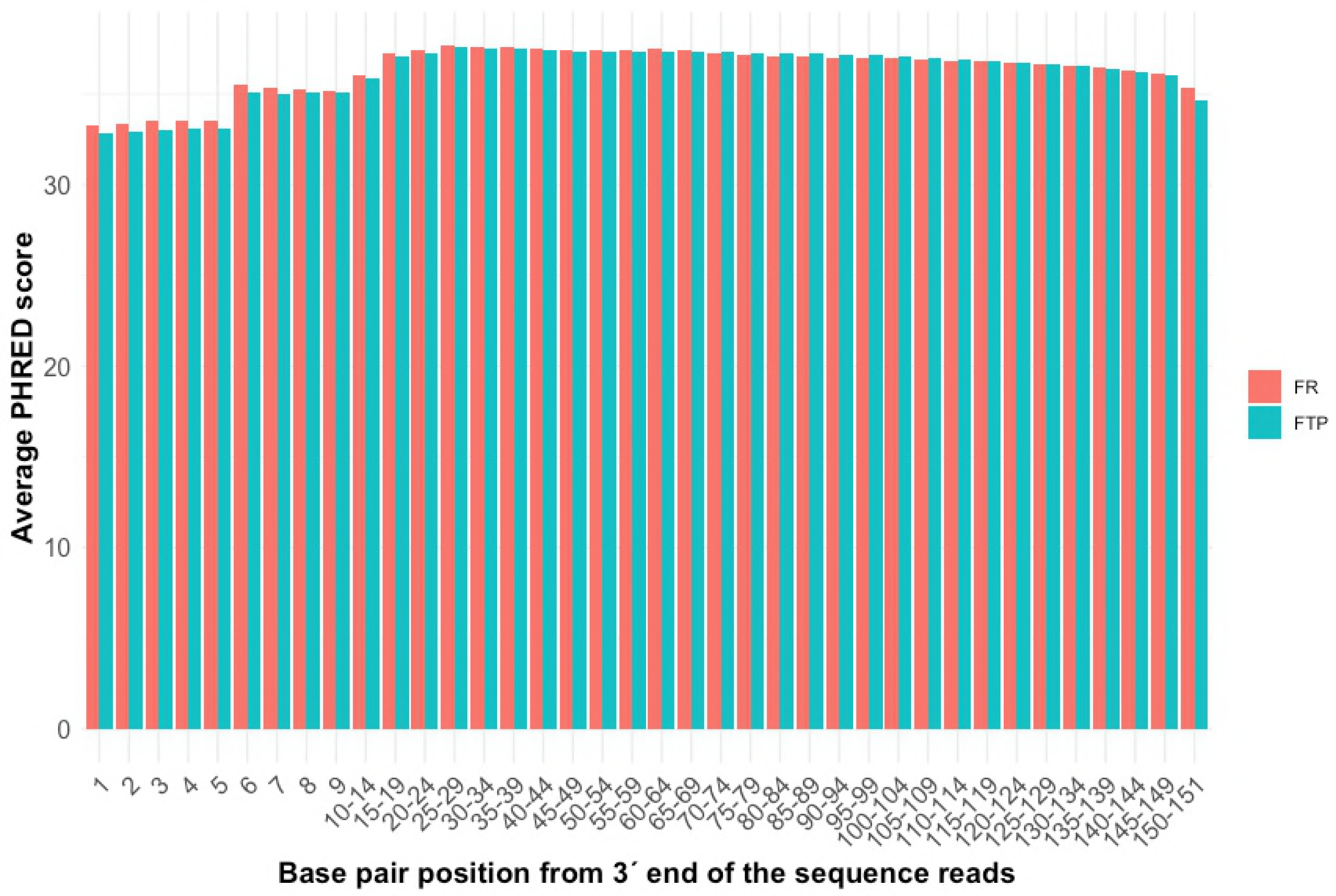
Comparison of the sequence qualities scores (PHRED) at the 38-ends of the sequences that have been generated from the NGS libraries constructed with Fragmentase (red) and FTP (blue) methods of DNA fragmentation. No difference was found between the libraries.

After NGS, the generated reads were subsequently aligned to the *E.coli* BL21(DE3) reference genome sequence (NCBI Ref Seq: NC_012971.2). There are several key characteristics of NGS that depend on a quality of the library: genome coverage, identity with a reference sequence, the rate of errors and amount of unmappable sequences. These characteristics were estimated for different sequencing depths of the NGS libraries. For the simulation of different depth, random samples of NGS reads were generated. To compare the genome coverage (the total number of aligned bases in the reference, divided by the genome size) we used the genome sequence NCBI Ref Seq: NC_012971.2 as the reference on the assumption this represented 100% coverage. Unmappable sequences were calculated as a rate of unmappable reads. A large fraction of these reads reduces the efficiency and the apparent coverage of the genome sequencing. The rate of indels was estimated as the average number of single nucleotide insertions or deletions per 100,000 aligned bases, and the rate of mismatches as the average number of mismatches per 100,000 aligned bases. The resulting average data of NGS analysis are shown in Table 1. The statistics for Fragmentase and FTP NGS libraries were calculated from the data of the four independent libraries for the each fragmentation method. The detailed data for each NGS library are shown in Supporting information (S1 Table). The obtained characteristics were about the same for the assembled sequences from the libraries generated by different methods (Table 1).

**Table 1.** **Key averaged NGS characteristics of Fragmentase and FTP generated libraries.**

| Sequencing depth<br>(numbers of reads) | Method of<br>DNA<br>fragmentation | Genome<br>coverage (%) | Ref. Seq.<br>identity (%) | Mismatch errors<br>(per 100 kb) | Indel errors<br>(per 100 kb) | Unmappable<br>reads (%) |
| --- | --- | --- | --- | --- | --- | --- |
| 32× depth<br>(10×10<sup>5</sup> reads) | Fragmentase | 98.226 | 99.999 | 1.01 | 0.24 | 3.07 |
|  | FTP | 98.224 | 99.999 | 1.02 | 0.14 | 3.91 |
| 16× depth<br>(5×10<sup>5</sup> reads) | Fragmentase | 98.193 | 99.999 | 1.05 | 0.13 | 3.09 |
|  | FTP | 98.200 | 99.999 | 1.17 | 0.16 | 3.92 |
| 8× depth<br>(2.5×10<sup>5</sup> reads) | Fragmentase | 98.042 | 99.996 | 3.70 | 0.22 | 3.17 |
|  | FTP | 98.068 | 99.996 | 4.02 | 0.24 | 3.90 |
| 3× depth<br>(1×10<sup>5</sup> reads) | Fragmentase | 91.100 | 99.974 | 25.23 | 0.70 | 3.13 |
|  | FTP | 90.908 | 99.971 | 27.70 | 1.21 | 3.90 |

The mean NGS statistics per library were calculated from the data of the four indepanded libraries for the each method. All metrics were obtained for different depths of *E.coli* BL21 genome sequencing. We found no significant differences between Fragmentase and FTP generated NGS libraries.

To evaluate the genome assembly *de novo* for Fragmentase and FTP libraries, we used QUAST software (quality assessment tool for genome assemblies) [9]. We compared the following assembling metrics:

- Number of contigs: The total number of contigs in the assembly.
- Largest contig: The length of the largest contig in the assembly.
- Total length: The total number of bases in the assembly.
- N50 and N75: The contig length such that using equal or longer length contigs produces at least 50% and 75% (accordingly) of the bases of the assembly length [9, 11, 12].
- NG50 and NG75 (Genome N50/75): The contig length such that using equal or longer length contigs produces at least 50% and 75% (accordingly) of the length of the reference genome, rather than 50% and 75% of the assembly length [9, 11, 12].

The assembly metrics were calculated for different sequencing depths of the libraries obtained by Fragmentase and FTP methods. The mean statistics calculated from the data of the four independent libraries for the each fragmentation method are shown in Table 2. The metrics for each NGS library are shown in Supporting information (S2 Table). Our results demonstrate that the characteristics of the genome assembly of libraries obtained by the novel FTP method are similar to those obtained by the Fragmentase method (Table 2).

**Table 2.** **The averaged assembly metrics of the NGS libraries obtained by Fragmentase and FTP methods.**

| Sequencing depth<br>(numbers of reads) | Method of DNA fragmentation | Number of contigs | Largest contig (bp) | Total length (bp) | N50 | NG50 | N75 | NG75 |
| --- | --- | --- | --- | --- | --- | --- | --- | --- |
| 32× depth<br>(10×10<sup>5</sup> reads) | Fragmentase | 182 | 272230 | 4485104 | 81481 | 80813 | 41851 | 40741 |
|  | FTP | 195 | 265892 | 4484951 | 81981 | 80479 | 43990 | 41542 |
| 16× depth<br>(5×10<sup>5</sup> reads) | Fragmentase | 204 | 217222 | 4483651 | 69766 | 69259 | 37530 | 36159 |
|  | FTP | 196 | 194626 | 4484098 | 70654 | 69018 | 39773 | 39008 |
| 8× depth<br>(2.5×10<sup>5</sup> reads) | Fragmentase | 304 | 134506 | 4478279 | 45010 | 44221 | 26078 | 24944 |
|  | FTP | 274 | 133551 | 4479908 | 41368 | 40106 | 21611 | 19769 |
| 3× depth<br>(1×10<sup>5</sup> reads) | Fragmentase | 2414 | 14250 | 4178082 | 2886 | 2689 | 1753 | 1476 |
|  | FTP | 2500 | 15256 | 4178040 | 2666 | 2456 | 1628 | 1348 |

The mean assembly statistics were calculated from the data of the four independent libraries for the each method and for the different depths of *E.coli* BL21 genome sequencing. No significant differences between Fragmentase and FTP generated NGS libraries were found.

In summary, the Fragmentation Through Polymerization is a novel, robust and simple method of DNA fragmentation, which is suitable for NGS. It simplifies the procedure and reduces the price of NGS library preparation by eliminating the DNA end-repair stage from the protocol. Thus, the FTP method can become a helpful tool for NGS.

## Acknowlegments

We thank Syntol JSC (Moscow, Russia), Evrogen JSC (Moscow, Russia) and Bioron GmbH (Ludwigshafen, Germany) for support of this project.

## Supporting information

**S1 Table. Key NGS characteristics of individual libraries generated by Fragmentase (A) and FTP (B) metods.** All metrics were obtained for different depths of *E.coli* BL21 genome sequencing.

**S2 Table. The assembly metrics of the individual NGS libraries obtained by Fragmentase and FTP methods.** All metrics were obtained for different depths of *E.coli* BL21 genome sequencing.

